# ZEB1 is Required for NHEJ-Mediated DSB Repair in Euchromatin

**DOI:** 10.1101/2020.05.13.094391

**Authors:** Thomas L. Genetta, Tarek Abbas, Raj Pandta, Clayton R. Hunt, Tej. K. Pandita, James M. Larner

**Affiliations:** Dept. of Radiation Oncology, University of Virginia School of Medicine, PO Box 800383, Charlottesville, VA 22908; Dept. of Biochemistry and Molecular Genetics University of Virginia School of Medicine, Charlottesville, VA 22908; The Houston Methodist Research Institute, Houston, TX 77030; Department of Molecular and Cellular Biology, Baylor College of Medicine, Houston, TX 77030

**Keywords:** ZEB1, 53BP1, NHEJ, euchromatin

## Abstract

Ionizing radiation-induced DSBs are repaired primarily by the Non-Homologous End Joining (NHEJ) pathway, but the details of how this is regulated in different chromatin contexts are far from understood. We have discovered a novel response to DSBs that promotes NHEJ selectively in euchromatin, based on a novel interaction between the EMT-inducing transcriptional repressor ZEB1, and the well-studied NHEJ-promoting DNA repair factor 53BP1. Using a number of approaches, we have discovered that the ZEB1-53BP1 association is amplified following exposure of cells to IR and that they co-localize at IR-induced foci (IRIF). Depletion of ZEB1 enhances radio-sensitivity and increases IR-induced chromosomal aberrations in an ATM-independent manner. The very rapid recruitment–within 2 seconds–of ZEB1 to euchromatic DSBs is like-wise ATM-independent, but DNA-PK-dependent and is required for subsequent recruitment of 53BP1. ZEB1 promotes NHEJ and inhibits HR through its homeodomain by inducing 53BP1-permissive, pro-NHEJ/anti-HR chromatin modifications. Lastly, depletion of ZEB1 increases hyper-resection at DSBs and inhibits physiological DSB repair. These results support the argument that ZEB1 plays an essential role in DSB repair in euchromatin by establishing a 53BP1-permissive/pro-NHEJ chromatin environment.

## Introduction

DNA double strand breaks (DSB) induced by ionizing radiation (IR), chemotherapeutic agents, or produced as by-products of physiological processes such as replication stress, transcriptional block or aberrant somatic recombination pose an existential threat to survival (1-3). Ionizing radiation induced DSB are mostly repaired by canonical Non-Homologous End Joining (cNHEJ) which is rapid and error prone. With the appearance of newly-replicated DNA in S phase and throughout G2, the predominant mode of DSB is by the error-free homologous recombination (HR) pathway. The details of how chromatin context, cell-cycle phase, cell type and differentiation state influence the DDR, regulating (among other things) the extent of strand resection and length of repair, are far from understood.

The **Z**inc finger **E**-box-**B**inding protein ZEB1 (also known as delta-EF1) has a well-documented role as a transcriptional regulator. ZEB1 and ZEB2 (SIP1) are the sole members of the zinc-finger-homeodomain family of transcription factors. Each has two clusters–amino and a carboxy-terminal–of C2H2-type zinc finger domains that can bind CANNT(G) elements (E-boxes) at promoters and enhancers and a central POU-like homeodomain, the function of which (in any context) has not been determined (4). Although initially described as transcriptional repressors through their interaction with the CtBP1/2 corepressors, ZEB factors can also activate transcription, through their interaction with coactivators (such as p300 and YAP (5, 6).

Through its activity at a number of genetic targets, notably the E-cadherin gene promoter, and at specific microRNA cluster loci, ZEB1 is one of a family of factors (incl. Snail, SLUG, TWIST, etc.) required for the initiation and maintenance of the epithelial-to-mesenchymal transition (EMT) during both development and tumor progression (7, 8). ZEB1 has been implicated in virtually all aspects of tumor biology, including initiation, progression, metastasis, cancer stem cell (CSC) maintenance (including induction of CSC-specific surface markers) and plasticity, and induces both chemo- and radio-resistance (9-16). Consistently, ZEB1 is highly expressed in both the interior of tumor masses, mirroring levels observed in relatively hypoxic stem cell (SC) compartments (as observed during normal development, tissue regeneration and tumor progression), and at the leading edge of invasive tumors (9, 11-13, 17). In the context of normal mammary duct development, high ZEB1 expression in epithelial stem cells is further implicated in suppressing oncogenic-induced genomic instability by up-regulating the methionine sulfoxide reductase anti-oxidant pathway (14). This affect appears to be tissue/tumor-type-specific, however, as ZEB1-mediated transcriptional repression of N-Methyl-Purine Glycosylase, a key enzyme initiating base-excision repair, contributes to inflammation-driven colorectal cancer (15). With respect to DSBs, recent work has shown that in response to IR, ATM phosphorylates and stabilizes ZEB1, which then recruits the deubiquitinase USP7 to stabilize CHK1, thereby extending the time available for DNA damage repair (18). Furthermore, it has been reported that ZEB1 transcriptionally regulates several DDR-related genes (19). Despite its potential importance to the DDR, however, essentially nothing is known about whether and how might ZEB1 function directly at DSBs.

Initially isolated based on its physical interaction with p53, 53BP1 is a key player in the DDR. 53BP1 rapidly localizes to DSBs, via recognition of specific ubiquitinated and methylated histone marks, to promote NHEJ by nucleating the anti-resection complex shieldin (20-25). 53BP1 is required for CSR, mid-range V[D]J recombination, and fusion of de-protected/dysfunctional telomeres (26). Depletion of 53BP1 radio-sensitizes normal as well as tumor cells in culture and in xenografts, dramatically increases the number and size of insertions and deletions (indels) at repair junctions, and increases chromosomal aberrations (27-31). While largely dispensable for cNHEJ in the context of therapeutically-induced DSBs, there is extensive evidence that 53BP1, via the shieldin complex, inhibits more error-prone NHEJ pathways (i.e. alternative end-joining and single-strand annealing) that rely on increasing levels of resection.

We present here data that demonstrate that ZEB1 is required for 53BP1-mediated effects on DSB repair in euchromatin. Our results demonstrate that 1) ZEB1 concentrates, in a DNA-PK-dependent manner, very rapidly at LASER-induced lesions, 2) establishes, through inhibition of both ATM and pro-HR chromatin modifiers, a pro-NHEJ environment, 3) physically interacts with and is required for the subsequent recruitment of 53BP1 to DSBs in euchromatin, further promoting NHEJ and inhibiting HR, and 4) is required for physiological distal NHEJ-mediated DSB end-joining. These data establish a novel link between ZEB1, a key mediator of the EMT, and 53BP1, a critical determinant of DSB repair by NHEJ.

## Results

### ZEB1 physically interacts with 53BP1 and they co-localize at DSBs

To identify interacting partners with which ZEB1 promotes therapeutic resistance we performed a yeast two-hybrid screen using fl human ZEB1 as the “bait”. Screening a HeLa cDNA library, 53BP1 was a prominent positively-interacting candidate. Reversing the configuration of the assay, with fl 53BP1 as the bait, the ZEB1 interaction (but not the negative controls) also resulted in colony growth. The interaction between 53BP1 and ZEB1 was confirmed in 293T cells via reciprocal co-immunoprecipitation (Fig 1A). In its role as a transcription factor, ZEB1 has been shown to nucleate a number of chromatin-modifying activities and co-factors, including the histone deacetylases (HDACs) 1 and 2 (32, 33), and both were detected in the ZEB1 IP (Fig. 1A), while only HDAC1 was detected in the 53BP1 IP (Fig 1B).

**Figure 1.**
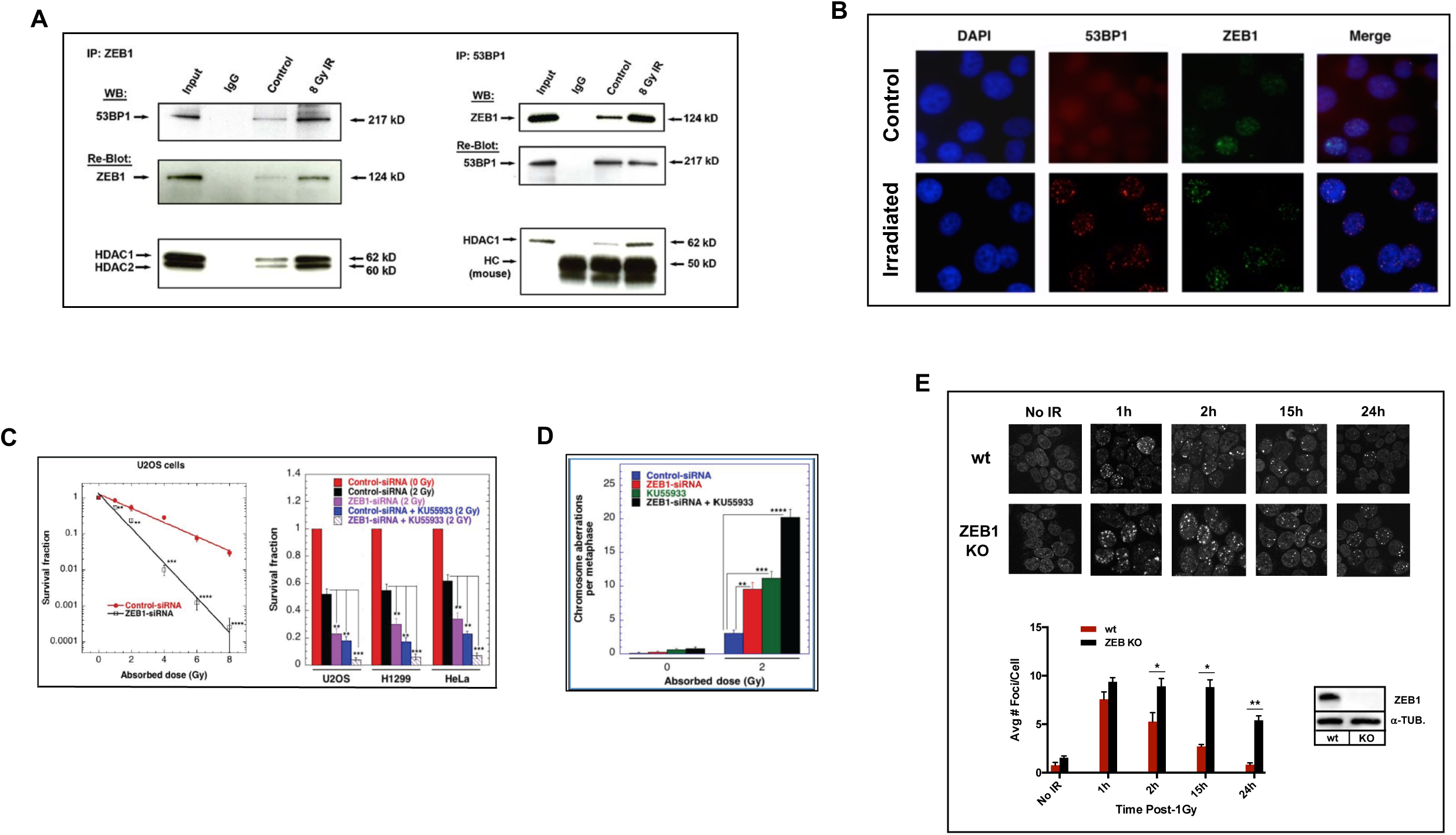
Endogenous interaction between ZEB1 and 53BP1 is amplified post-IR. (A) Reciprocal IP, followed by representative WB, shows that in U2OS cells, ZEB1 or 53BP1 each co-IP the other, as well as HDAC1/2, an interaction that is amplified 30 min after 8 Gy IR; HC = Ig heavy chain. (B) IF staining of 293T cells shows that ZEB1 and 53BP1 co-localize at IRIF. Red, 53BP1; Green, ZEB1; blue, DAPI; Top: No IR control; Bottom: 30 min. post-6Gy. (C) ZEB1 KD increases radio-sensitivity, independent of ATM. Left panel, Clonogenic survival of U2OS cells transfected with anti-ZEB1 or NT siRNA; Right panel, Clonogenic survival of three cell types, post-2Gy, +/- ATM inhibitor KU55933, +/- ZEB1 KD; (D) ZEB1 KD increases chromosomal aberrations independent of ATM. 293T cells transfected with either anti-ZEB1, with or w/o KU55933, or non-targeting siRNA and 48h later subjected to 2Gy IR. Colcemid was added 5h post-IR, and metaphase spreads prepared as described (X); (E) ZEB1 KO delays resolution of gH2AX foci. Top: IF staining time course of appearance, post-1Gy, and resolution of gH2AX foci, in either wt (top row), or ZEB1-depleted (via CRISPR KO) 293T cells visualized via IF staining, Bottom, left: quantitation of staining results; Bottom right: WB shows virtual absence of ZEB1 protein in the KO line. Data shown in all panels is a representative example of a minimum of three independent experiments. Asterisks indicate statistically significant differences by the Student’s *t* test; NS, no significance, *P* > 0.05; \**P* ≤ 0.05, \*\**P* ≤ 0.01, \*\*\**P* ≤ 0.005, \*\*\*\**P* ≤ 0.001.

Interestingly, we observed that the relative amounts of both ZEB1 and 53BP1 in the co-IP increased 30 min post-IR (Fig. 1, lane 4 in each panel). Double immunofluorescence staining of 293T cells exposed to 6 Gy showed endogenous co-localization of ZEB and 53BP1 to IR-induced foci (Fig 1B).

### ZEB1 depletion increases radio-sensitivity and chromosomal aberrations, independent of ATM

ZEB1 is a well-characterized driver of both radio-and chemo-therapeutic resistance (9, 11) and has recently been shown to mediate radio-resistance in a breast cancer cell line, in part, via stabilization of CHK1 kinase, an effect that is ATM-dependent (13). As expected, ZEB1-depleted U2OS, HeLa, and H1299 cell lines, sustained increased IR-induced cell killing, an effect nearly as strong as inhibition of ATM. Interestingly, in all three cell lines, sensitivity was further augmented by combining these two treatments (Fig. 1C, right panel), suggesting that ZEB1 engages a pathway distinct from ATM in response to IR.

Analysis of metaphase spreads revealed that ZEB1 depleted cells have a 3-fold increase in chromosome aberrations, including breaks, gaps, radials and translocations. This effect was nearly doubled when IR and pharmacological inhibition of ATM were combined (Fig 1D). Thus, ZEB1 protects cells from the effects of IR and limits chromosome instability following IR.

### Loss of ZEB1 Delays Resolution of γH2AX foci

Wild-type or Crisper/Cas9-generated ZEB1 KO U2OS cells were subjected to 1Gy, and appearance and resolution of γH2AX foci, a surrogated marker of DSB, were monitored over the next 24hrs. Residual γH2AX foci were significantly greater in the ZEB1 KO cells compared to control (Fig. 1E, quantitated in lower panel).

### ZEB1 levels increase 4-fold at DSBs in euchromatin, independent of ATM

Given that ZEB1 and 53BP1 physically interact, that they co-localize at DSBs, and that ZEB1 is required for resolution of γH2AX foci, we next sought to determine the extent to which these two factors regulate each other at a “clean” enzymatically-induced DSB, and whether this is affected by the chromatin environment/conformation. To answer these questions, we employed our previously characterized 293T-based stable cell lines that harbor CRISPR-Cas9-knocked-in homing endonuclease I-SceI target sites into two euchromatic and two heterochromatic regions, in this case, on chromosome 1. As demonstrated in previously, we verified, for each of these four sites, that they a) retained relevant chromatin conformation-specific epigenetic marks, b) that they can each be efficiently cleaved by a transfected cDNA encoding the I-SceI mega-nuclease, and c) chromatin context does not affect repair efficiency (34, 35). We used this system first, to evaluate the relative levels of ZEB1 and 53BP1 before and after induction of DSBs. Baseline (pre-cleavage) levels of ZEB1 in the two euchromatic regions (Fig 3A, regions A and D) far exceed those in the 2 heterochromatic regions (regions B and C) as determined by ChIP/qPCR. Strikingly, introduction of a DSB (via transfection of an I-SceI expression vector) induced an approximately 4-fold increase in ZEB1 levels in the euchromatic regions, with virtually no change in levels at the heterochromatic sites (Fig 3A). Using siRNA KD, we further demonstrated that this DSB-mediated increase in ZEB1 levels was independent ATM function (Fig 4B). Our findings that KD of ZEB1 results in an IR-induced decrease in survival, an increase in chromosomal aberrations, and reduction in 53BP1 localization to “clean” (enzymatically-induced) DSBs suggest that ZEB1 functions at DSBs independently of the ATM-USP7-CHK1 pathway (17).

**Figure 2.**
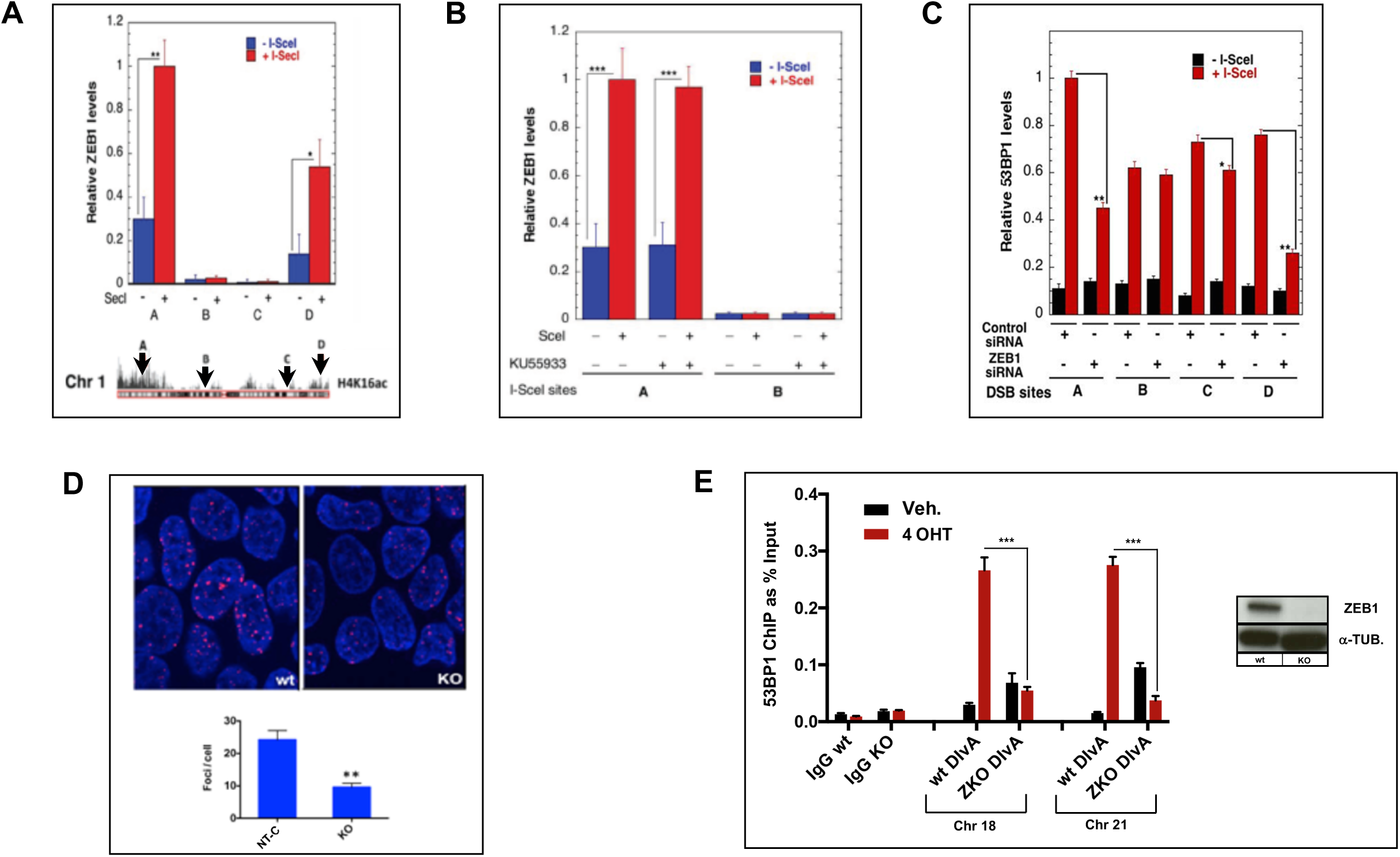
ZEB1 is required for localization of 53BP1 to enzymatically-induced euchromatic DSBs. (A) ZEB1 levels increase 4-fold at DSBs in euchromatin, independent of ATM. Top: Following induction of an enzymatic DSB in each of four 293T-based stable lines–sites denoted A-D in lower schematic–cells were processed for ChIP to detect ZEB1 binding near the cut; Bottom: Schematic of chrom. 1 with CRISPR-Knocked-In I-SceI sites, two (A & D) in H4Ac16-rich (euchromatic) and two (B & C) in H4Ac16-poor (hetero-chromatic) regions; (B) Same expt. protocol as in panel A, showing siRNA KD of ATM has no effect on the increase in ZEB1 levels post-DSB at I-Sce I site A; (C) ZEB1 loss impairs recruitment of 53BP1 to DSBs. Forty-eight hrs post-treatment with either control or ZEB1-targeted siRNA, DSBs were induced in each of the four stable cell lines in depicted in panel A, processed and then analyzed by ChIP/qPCR. Relative levels of 53BP1 at euchromatic DSB sites A & D are significantly reduced with ZEB1 KD, with no or lesser effect at their heterochromatic counterparts. (D) Top, representative images of wt or CRISPR ZEB1 KO 293T (see Fig. 1E for WB) subjected to 6 Gy, fixed 20min later and stained for 53BP1 via IF; below irradiated ZEB1 KO cells have fewer than half the 53BP1-IRIF compared with wt control; (E) Loss of ZEB1 dramatically reduces localization of 53BP1 800 bp from AsiSI-induced DSBs. ChIP/qPCR analysis of 53BP1 binding near AsiSI-generated DSBs on either chromosome 18 or 21, in wt or ZEB1 KO DIvA U2OS cells. IgG neg. control was carried out using the chr. 18 primer set. Right panel is a representative WB showing absence of ZEB1 protein in the CRISPR KO DIvA cell line. In panels A and B, C and E, bar graphs depict relative ChIP/qPCR results, adjusted for background (ChIP with non-specific IgG) and normalized to Input. IF staining analysis of ZEB1 effects on IRIF and all ChIP/qPCR experiments were carried out at least three times. NS, no significance, *P* > 0.05; \**P* ≤ 0.05, \*\**P* ≤ 0.01, \*\*\**P* ≤ 0.005.

**Figure 3.**
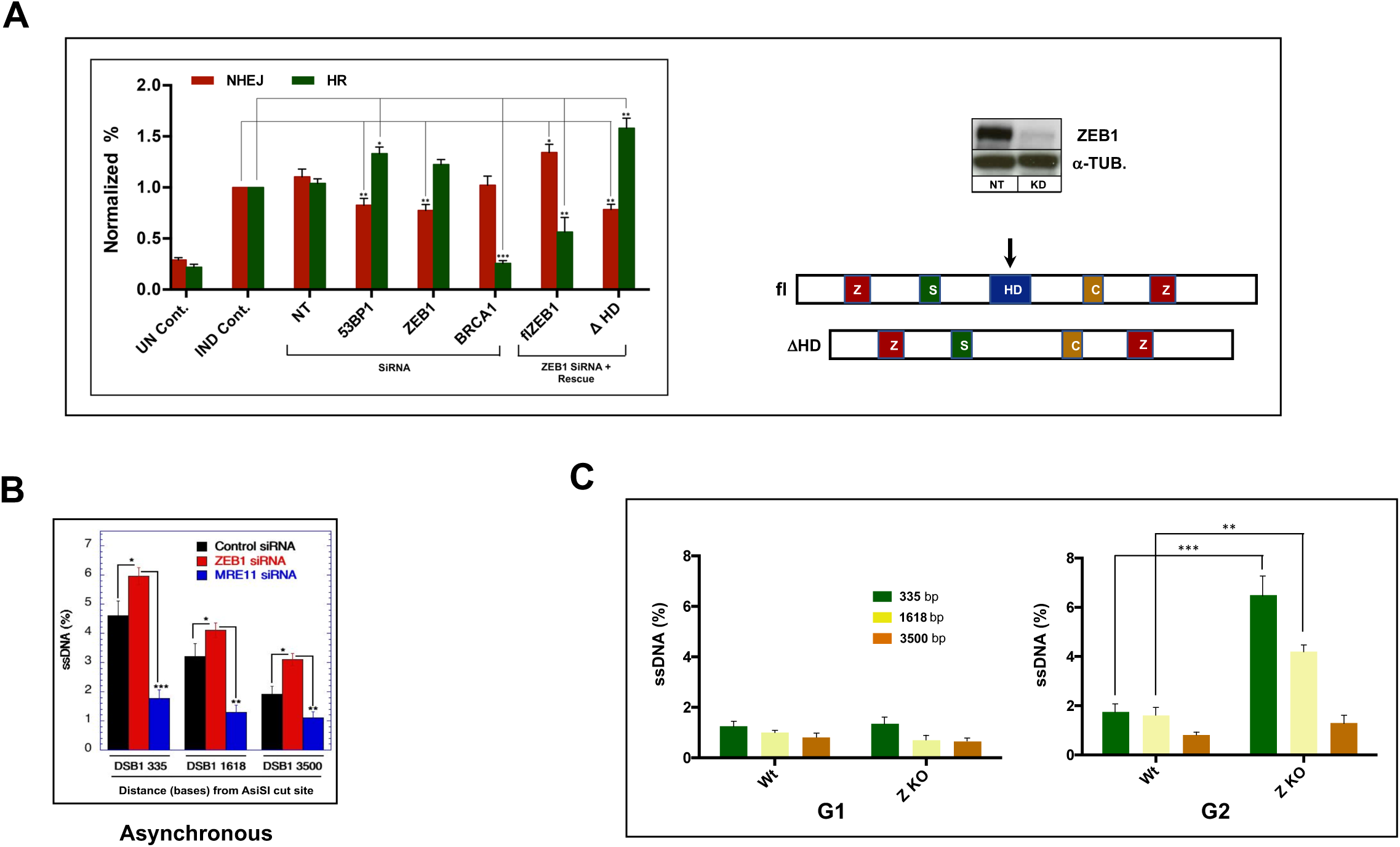
ZEB1 promotes error-prone NHEJ to a similar extent as 53BP1 and attenuates HR. The EJ-DR U2OS-based cell line with stably integrated, I-Sce I-based reporters for both error-prone NHEJ (RFP) and HR (GFP), were transfected with either NT or indicated SiRNAs, and 72h later, DSBs were induced (see Methods; X, Y). Cells were subjected to FLOW analysis 72h after DSB induction. The anti-ZEB1 siRNA targeted its 3’UTR, allowing rescue by either full-length (fl) or a homeodomain-deleted (ΔHD) cDNAs (see schematic, right panel). ZEB1 KD, similar to the effect of 53BP1 KD, results in 20% reduction in error-prone repair, and more modest effect on HR. This effect is rescued, with a potentiation of NHEJ and a concomitant reduction in HR upon transfection of the fl ZEB1 expression vector. Transfection of the ΔHD construct had the opposite effect. Avg of three independent experiments with SEM, normalized to induced (IND) control; UN Cont., un-induced/No SiRNA; IND. Cont., induced/No SiRNA. error bars = SEM; right panel, above is a representative WB showing the effect of the anti-ZEB1 siRNA; in schematic, ZF, Zinc Fingers; SID, SMAD-Interacting Domain; HD, Homeodomain; CID, Co-Repressor-Interacting Domain. (B,C) ZEB1 Inhibits Resection Following an Enzymatic DSB in. AsiSI-generated DSBs were induced in (B) asynchronous or (C) synchronized DIvA cells (see Fig. 1E for WB of ZEB KO cells), genomic DNA isolated, digested with restriction enzyme BsrGI and subjected to qPCR (in triplicate) using PCR primers that flank BsrGI sites at 335pb, 1618bp or 3500 bp downstream of an AsiSI cut site on chromosome I (X). Resection, yielding increased levels of SS DNA and subsequent PCR product, is increased in asynchronized cells with ZEB1 KD, (with MRE11 KD as a positive control), a result amplified in G2 (C). Results, normalized to AsiSI cutting efficiency, are the avg. of three different expts. with SEM. *P* > 0.05; \**P* ≤ 0.05, \*\**P* ≤ 0.01, \*\*\**P* ≤ 0.005.

**Figure 4.**
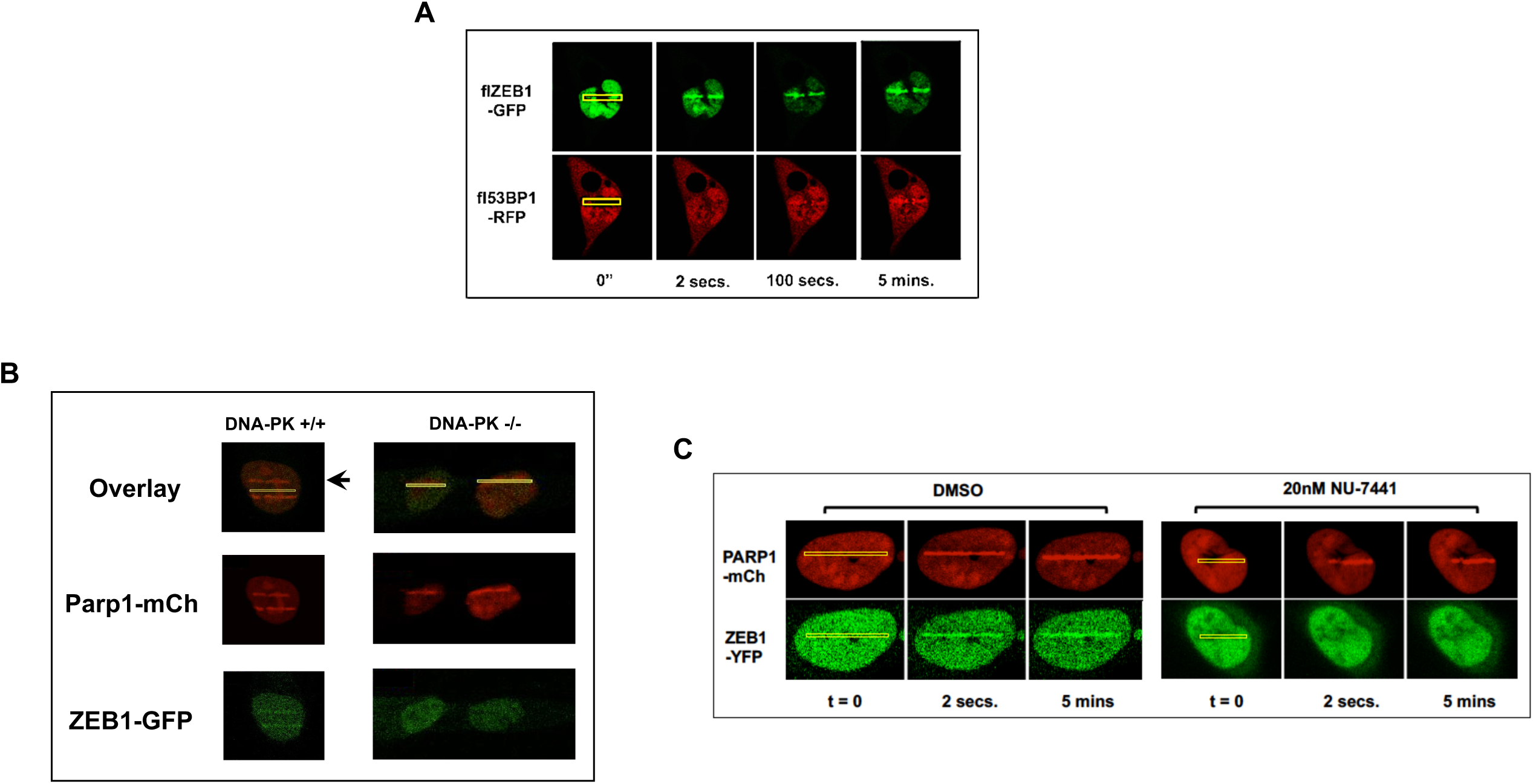
ZEB1 localization to LASER-induced Lesion is rapid, precedes 53BP1 and requires DNA-PK activity. (A) U2OS cells transfected with the indicated fluorescence-tagged cDNAs and 48h later were subjected to 780nM Near-Infrared LASER ablation in the stripes defined by the yellow rectangles. Frames were continuously captured (2 secs total for both channels), images extracted. ZEB1-GFP concentrates at the lesion within 2 secs.; 53BP1-RFP co-localizes about 1.5 mins later. (B) The GBM cell lines M059K (DNA-PK^+^) and M059J (DNA-PK -) were transfected with the indicated fluorescence-tagged cDNAs and 48h later LASER-mediated lesions were induced as in panel (A). In the absence of DNA-PK, ZEB1 fails to localize to the lesion. Arrow in panel B indicates a lesion induced 5’ prior to the continuous capture depicted here, suggesting that the ZEB1-GFP/PARP1-RFP-decorated stripe persists for at least that long after the advent of a DSB. (C) ZEB1 localization is eliminated with DNA-PK inhibitor NU 7441 (20nM, 1hr prior to ablation). In panels B and C, transfected PARP1-RFP serves as a relative temporal marker. Captured images are representative of at least three separate transfections/analyses (done on different days).

### ZEB1 depletion impairs 53BP1 recruitment to DSBs in euchromatic regions

We next asked whether recruitment of ZEB1 to euchromatic DSBs affected 53BP1 localization. Strikingly, at both euchromatic I-SceI-mediated DSBs, ZEB1 KD significantly impaired 53BP1 enrichment (via ChIP/qPCR), with a minor effect at one of the DSBs in a heterochromatic region (Figure 2C). Employing a CRISPR/Cas9-mediated ZEB1 KO 293T cell line, we next obtained a visual corroboration of the reduction in 53BP1 localization to DSBs. Wild-type or ZEB1 KO cells were irradiated with 6Gy, and after 20 min, fixed and stained for endogenous 53BP1 via IF. Compared with wt, ZEB1 KO cells have approximately half the 53BP1 foci (Fig 2 D; quantified below, left).

The ChIP result was reproduced using the U2OS-based (p53 wt), DIvA (**D**ouble-stranded breaks **I**nduced **v**ia **A**siSI) cell line, a 4-OHT-inducible system, that allows for the reliable interrogation of approximately 100 AsiSI endonuclease-targeted sites dispersed throughout the human genome (virtually all of these reside in euchromatin, previously mapped and characterized, (36, 37). This number of DSBs provides a reasonable, enzymatically-generated analogue to the cellular consequences of a therapeutic dose of IR (36, 38, 39). As above, we generated a ZEB1 KO derivative of this line and, following induction of DSBs, determined, via ChIP/qPCR, the extent of 53BP1 localization to two sites, comparing the parent line to ZEB1 KO. The reduction in 53BP1 localization at induced DSBs in chromosomes 18 and 21 in the ZEB1 KO DIvA line exceeded the levels seen in Fig 2C with ZEB1 KD, (Fig. 2E).

### ZEB1 promotes error-prone NHEJ to a similar extent as 53BP1 and attenuates HR

To determine whether ZEB1 influences NHEJ or HR pathways (or both) we utilized the EJ-DR U2OS cell line, which has three stably-integrated constructs: reporters measuring either error-prone NHEJ- (RFP) or HR- (GFP) mediated repair, and a cDNA encoding an inducible I-SceI enzyme (40). Fluorescence output is dependent on successful, pathway-specific repair (as opposed to unrelated or off-target effects). Cells were transfected with siRNAs targeting ZEB1, 53BP1, BRCA1, or NT control, and, following induction of the I-SceI enzyme, processed for analysis by FLOW (40, 41). The results, normalized to controls, clearly show that ZEB1 promotes error-prone NHEJ to a similar extent as 53BP1 (Fig 3A).

### ZEB1 Inhibits Resection Following an Enzymatic DSB in G2-arrested cells

Using asynchronous DIvA cells, and a qPCR-based resection assay (42), we further found that ZEB1 inhibits HR, at least in part, through the attenuation of resection activity (Figure 3B). This effect was virtually all confined to the G2 phase as shown in double-thymidine-blocked ZEB1 KO DIvA cells (Figure 3C). As ZEB1 is required for localization of 53BP1 to enzymatically-induced DSBs in euchromatin (Fig 2C-E) this result is consistent with the hypothesis that it is required for 53BP1/shieldin-mediated-block to resection activity (20-25), and ZEB1 depletion significantly increased that activity up to at least 3500 bases from the AsiSI-mediated DSB (Figs 3A,B).

### Recruitment of ZEB1 to LASER-induced lesions is DNA-PK-activity-dependent

We next sought to determine whether both factors arrived coincidentally to a DSB, or if ZEB1 preceded 53BP1. Using continuous-capture, live cell imaging and near-infrared LASER ablation, we introduced a complex lesion (including DSBs, with a minimum of pyrimidine dimers, (43) into nuclei of U2OS cells co-expressing ZEB1-GFP and 53BP1-RFP. We found that localization of ZEB to the lesion was very-rapid–within at least 2 seconds of the introduction of the lesion–and precedes 53BP1, which begins to decorate this lesion about 90 seconds post-irradiation (Fig 4A).

As we had demonstrated that ZEB1 localization to enzymatically-induced DSBs was ATM-independent, we next asked whether the DSB-triggered apical kinase DNA-PK, a critical component for cNHEJ, might play a role. To determine whether absence of DNA-PK affected ZEB1 localization to a LASER-induced lesion, we employed the isogenic glioblastoma cell lines, MOK (DNA-PK +/+) and MOJ (DNA-PK -/-; (44). Because of its well-characterized, very rapid DSB-localization kinetics (45), we used PARP1-RFP as relative timing factor in these analyses. Absence of active DNA-PK in the MOJ cells, eliminated the ZEB1-GFP-decorated lesion seen within the 2 second time-frame in the MOK line (Fig 4B). We then tested whether loss DNA-PK activity was responsible for this effect. Strikingly, ZEB1-GFP localization was eliminated in the presence of 20nM of the DNA-PK inhibitor NU-7441 (Fig 4B). We next investigated the contribution of ZEB1 in establishing a pro-NHEJ/anti-HR chromatin environment.

### ZEB1 inhibits HR-promoting Chromatin Modifying Activities

We have established that recruitment of pro-NHEJ/anti-HR factor 53BP1 to an enzymatically-induced DSB in euchromatin requires prior ZEB1 recruitment (Fig 2C,D,E) and that the latter event is dependent on DNA-PK activity (Fig 4B). In this regard, ZEB1’s role in transcription as a nucleator of chromatin modifying activities could reasonably be assumed to have been conscripted by the DDR machinery to help establish a pro-NHEJ/anti-HR chromatin environment. Initiation of the HR pathway includes the ATM-dependent rapid recruitment of methyl-transferase activities to increase the levels of H3K9me2/3, establishing an anti-53BP1/pro-BRCA1 chromatin environment (46, 47). We next asked whether loss of ZEB1 affected the status of H3K9me2/3 and observed that ZEB1 depletion in the I-SceI-inducible 293T cell line led to a significant increase in H3K9me2/3 levels (Fig 5A). Furthermore, using the ZEB1 KO DIvA cells, we detected over 10-fold increases (compared to controls) in two key components of the H3K9me2/3-inducing, chromatin condensing, pro-HR module that includes the methyl-transferase PRDM2 and its localizing partner, histone variant macroH2A1 (Fig 5B,C; (47). These results support the argument that rapid recruitment of ZEB1 to euchromatic DSBs is required for the establishment of pro-53BP1/NHEJ chromatin remodeling (summarized in Fig 5D).

**Figure 5.**
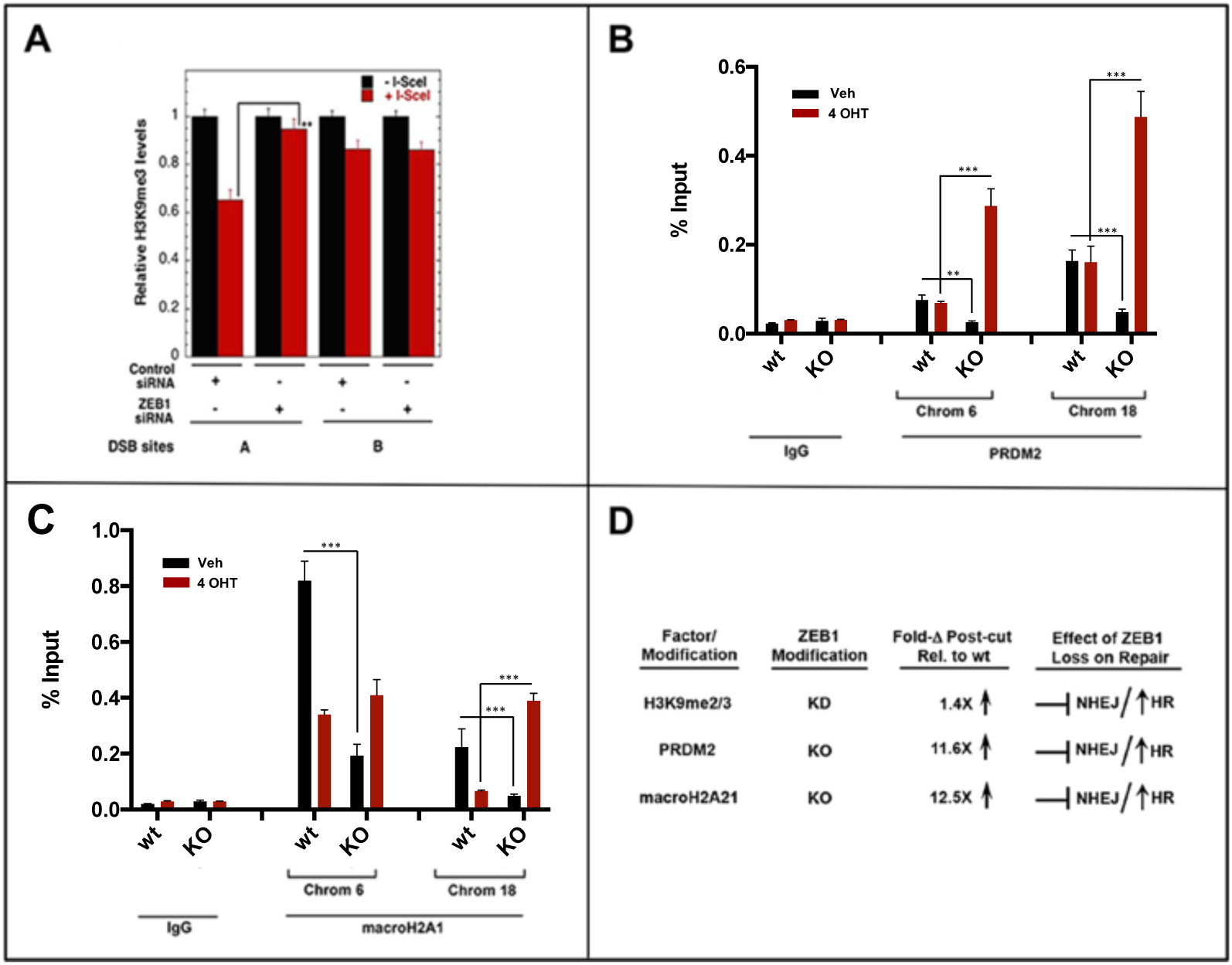
ZEB1 inhibits HR-promoting activities, promoting NHEJ. (A) Following I-SceI-mediated DSB induction (293T cells), ZEB1 KD increases the HR-promoting H3K9me3 modification, assessed via CHIP/qPCR; (B), (C) localization of the H3K9me2/3- and HR-promoting factors, macroH2A1 and the methyl-transferase PRDM2 are greatly increased in the absence of ZEB1 at two distinct DSBs in the U2OS-based DiVA cell line 4h post-induction (see WB in Fig 2E for ZEB1 KO DIvA cells); (D) summary of these effects, which together strongly suggest that ZEB1 acts to inhibit HR-promoting chromatin modifiers to promote NHEJ. In panels A, B, and C, bar graphs depict relative ChIP/qPCR results, adjusted for background (ChIP with non-specific IgG) and normalized to Input. All ChIP/qPCR experiments were carried out at least three times. NS, no significance, *P* > 0.05; \**P* ≤ 0.05, \*\**P* ≤ 0.01, \*\*\**P* ≤ 0.005.

### ZEB1 Inhibits HR Thorough its Homeodomain

Several very recent reports have begun to illuminate the distinct roles of ATM vs DNA-PK in DSB repair. These two DDR apical kinases can cross-regulate each other both positively and negatively, depending on the complexity and chromatin context of a DSB. Each can promote repair in specific chromatin contexts (48) and, at euchromatic, AsiSI-induced, “clean” DSBs, DNA-PK can inhibit ATM via phosphorylation, as well as resection (49, 50), promoting cNHEJ and inhibiting HR, and ATM promotes HR-permissive chromatin remodeling (47). We have shown above that rapid localization of ZEB1 to a LASER induced lesion requires DNA-PK kinase activity and that ZEB1 is itself required for 53BP1 recruitment to and inhibition of resection. Interestingly, in addition to its role as a localizer of chromatin modifying activities, a structural aspect of ZEB1 protein–the presence of a homeodomain (4); see schematic in Fig 3A)– connects these findings in a fundamental way. As homeodomain-containing transcription factors have been shown to regulate ATM activity (both positively and negatively; (51, 52)), we were prompted to ask whether this property of ZEB1 could also play a role in promoting the pro-DNA-PK/NHEJ-anti-ATM/HR repair outcome. Depletion of ZEB1 in the EJ-DR reporter cell line reduced NHEJ-mediated fluorescence and up-regulated the HR fluorescence output (Figure 3A). Following rescue of this phenotype by introduction of a full-length ZEB1 cDNA, this fluorescence readout was rescued, supporting a role for ZEB1 in the promotion of NHEJ and inhibition of HR. Interestingly, introducing a ZEB1 cDNA with only the Homeodomain region deleted (ΔHD, Figure 3A) failed to rescue this repair phenotype. This result, combined with the requirement for DNA-PK for ZEB1 localization/stabilization at LASER-induced lesions (Fig 4B), implicates the cooperation of these two factors in the earliest stages of the determination of DSB repair pathway choice, promoting NHEJ at the expense of ATM-initiated HR-permissive chromatin remodeling.

### ZEB1 is required for physiological DSB repair

Given its DNA-PK-dependent very early timing of localization–well before 53BP1 recruitment–we asked whether ZEB1 might play a role in physiological cNHEJ, cooperating with DNA-PK to promote rapid end joining at enzymatically induced DSBs that require minimal processing. Extensive characterization of Class Switch (CSR) and Variable [Diversity] Joining (V[D]J) recombination have demonstrated an absolute requirement for 53BP1 in the former (53-55) and for a subset of recombination events in the latter (56). Both of these programmed, distal end-joining pathways also have strict requirement for DNA-PK activity for repair of their respective, enzymatically-induced DSBs (57, 58). A role for ZEB1 in V[D]J recombination is further supported by a phenotypic outcome in a ZEB1 loss-of-function mutant mouse, which harbors a 99% reduction in the total thymic T cell population (59). While the spleens of these animals appear normal, an analysis of the B cell compartment at the level of class-switching (to our knowledge) has not been conducted (see discussion). We therefore asked, as is the case for non-physiological DSBs, whether ZEB1 might play a role in normal somatic recombination. To investigate CSR, we employed the standard CH12.F3 mouse B cell line, which, after challenge with a cocktail of TGF-b, IL-4 and anti-CD40 Ab, carries out an enzymatically-induced (via Activation-Induced Deaminase) distal NHEJ reaction in the constant region of the immunoglobulin heavy chain gene, resulting in a fusion of the previously recombined V[D]J region from the “M” to the “A” isotype, which is detected on the cell-surface via FLOW (60). Depleting ZEB1 using specific siRNA resulted in a greater than 50% reduction in cell-surface expression of IgA (Fig 6A).

**Figure 6.**
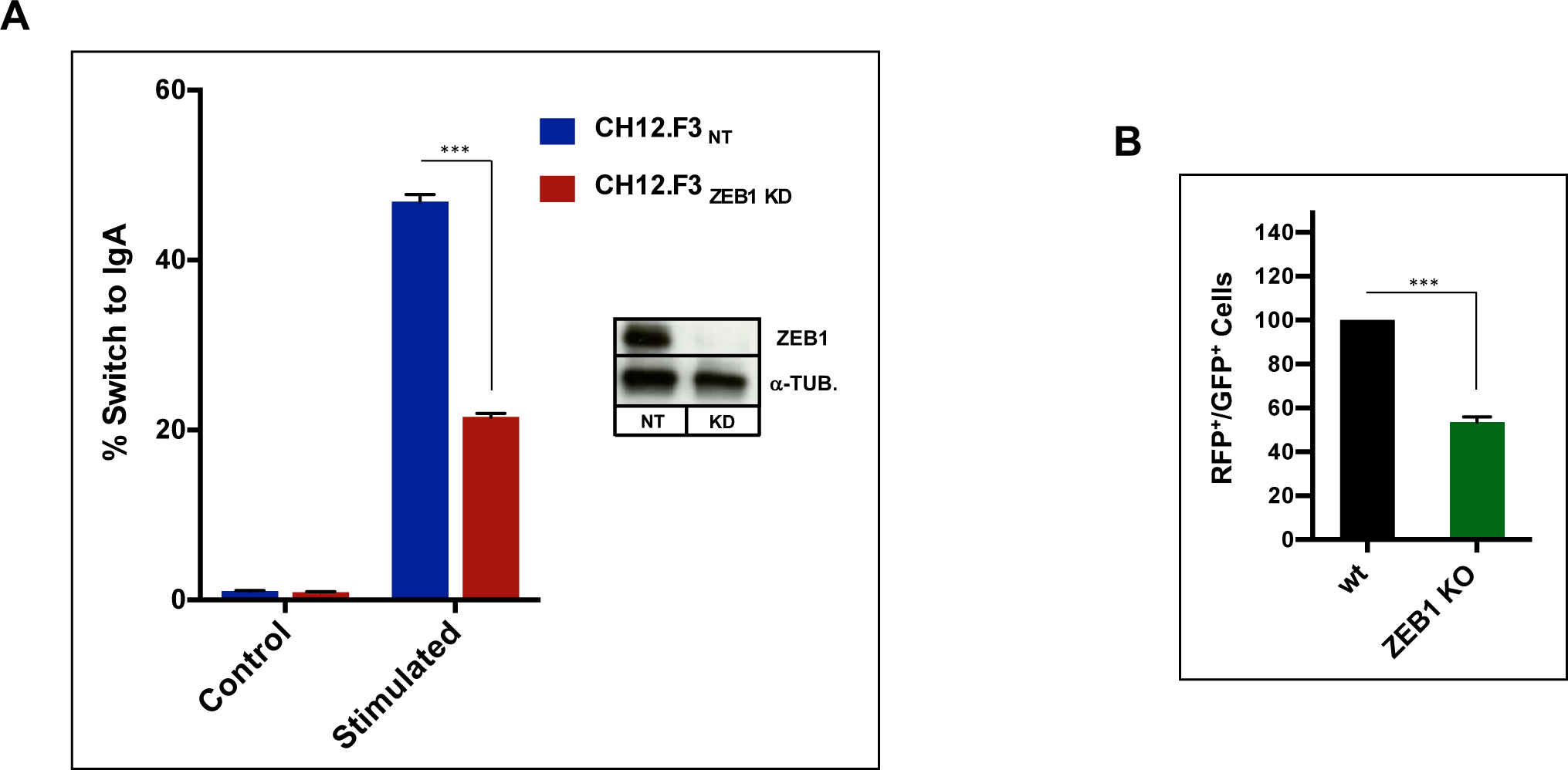
ZEB1 is required for physiological DSB repair. (A) Class Switch recombination: the murine B lymphocyte cell line CH12.F3 was electroporated with either non-targeting (NT) or anti-ZEB1 SiRNA (ZEB1 KD) and 48h later stimulated to undergo CSR. After 72h cells were stained and gated live cells (7AAD negative) analyzed by FLOW for % of surface IgA expression. ZEB1 KD resulted in greater than 50% reduction in IgA-positive cells. (B) V[D]J recombination: wt or ZEB1 KO 293T cells (see Fig. 1E for WB) were transfected with the RAG-responsive GFPi (inverted, non-expressed GFP cDNA) reporter and CMV-driven expression vectors for RAG1 and RAG2 and gated live cells (7AAD negative) scored 48h later for percentage of GFP-positive signal (inside an RFP+ gate – see methods). Loss of ZEB1 resulted in greater than 50% reduction in recombination-mediated inversion of the GFP cDNA to the expressed orientation; \*\*\**P* ≤ 0.005.

To investigate the role of ZEB1 in V[D]J recombination (in the context of 53BP1 involvement, the result of long-range distal end-joining of two enzymatically-induced DSBs; (56), we made use of a straight-forward fluorescent reporter-based test system. Co-transfection with cDNAs expressing the Recombination Activation Genes, RAG1 and RAG2, results in either an NHEJ-mediated inversion of a GFP cDNA into the correct orientation for expression downstream of a constitutively acting promoter or as an excised, non-expressing fused ring (61). When we introduced these three constructs into ZEB1 KO 293T cells, we observed, via FLOW, an approximately 40% reduction in the establishment of productive, GFP-expressing cells compared to wt controls (Fig 6B). Taking into account that others have shown the distance between the distal DSBs is determinative for the requirement for 53BP1 in V[D]J (56) this result is suggestive that ZEB1 is required for at least a subset of physiological distal NHEJ repair.

## Discussion

A well-characterized transcription factor, ZEB1 is known for promoting the EMT in both development and in metastasis, as well as for having pleiotropic effects that collectively potentiate tumor progression (8, 17). Given that ZEB1s well-established function is to nucleate chromatin modifying/re-modeling factors/activities at promoters and enhancers (32, 62), it’s rapid localization–within two seconds–to the sites of DSBs, is consistent with ZEB1 playing a similar role in DSB repair. The data presented here strongly suggest that ZEB1 acts in cis at DSBs in euchromatic regions and promotes the establishment of an anti-HR, pro-53BP1-premissive chromatin environment. We note that our data do not rule out the possibility that ZEB1 may also transcriptionally regulate genes involved in the DDR (19).

It has been reported that, in response to IR exposure, ATM phosphorylates and stabilizes ZEB1, which then recruits the deubiquitinase USP7 to stabilize CHK1, thereby extending the time available for DNA damage repair (17). In addition, it shows that ZEB1 promotes HR repair in a radio-resistant derivative of the human breast cancer HMLER cell line (63). In contrast, our data, derived using a number of different approaches, including an I-SceI Knock-In 293T cell line, the AsiSI U2OS DIvA cell line, live cell imaging, the EJ-DR pathway-choice reporter system, and CSR and V[D]J recombination reporters, collectively demonstrate that in euchromatin, ZEB1 is required for 53BP1 localization, NHEJ-mediated repair, and inhibits an HR-permissive chromatin environment. Though we cannot reconcile these conflicting out-comes, it is possible that the increase in ZEB1 protein levels seen in the radio-resistant variant of the HMLER cell line results in a titrating/squelching effect on NHEJ-related pathway components, such as 53BP1, reducing their ability to act in cis at a DSB, and allowing strand resection.

From a mechanistic standpoint, we show that ZEB1 promotes an NHEJ-permissive chromatin environment, in part by inhibiting the macroH2A1.2-PRDM2 methyltransferase complex, which deposits the pro-HR H3K9me2/3 mark. The availability of the HR pathway (and strand-resection) is up-regulated in S-phase to maximize error-free repair (64). A large body of work over the last decade has begun to clarify the earliest remodeling events that participate in establishing an HR-permissive chromatin environment. Briefly, results from LASER ablation and also biochemical studies, suggest that in the immediate aftermath of a euchromatic DSB, adjacent chromatin undergoes a fleeting expansion, followed by a rapid contraction. This has been interpreted as remodeling activities first clearing the area around the break (e.g. expulsion of nucleosomes and transcriptional machinery) and then imposing a BRCA1-permissive, nucleosomal/epigenetic environment (47, 65, 66). In that latter contraction phase, ATM-dependent recruitment of the histone variant macroH2A1 and the methyltransferase PRDM2 results in establishment of the H3K9me2/3 chromatin-contracting mark (46, 47). We report here that, in the absence of ZEB1, these three key chromatin-contracting elements–macroH2A.1, PRDM2, and H3K9me2/3–are all significantly increased, suggesting that one role of ZEB1 is to counteract the establishment of a BRCA1-permissive environment; the near-immediate localization of ZEB1 to a LASER lesion further supports this. Such activity is further supported by our finding that ZEB1s homeodomain is required for the pro-NHEJ/anti-HR fluorescence output of the EJ-DR pathway choice cell line (Fig 3A), suggesting that, like many homeodomain-harboring proteins, ZEB1 may serve as a block to ATM activity in this context, favoring DNA-PK-mediated direct end-joining over DSB repair by the HR pathway (52).

The mechanistic role for 53BP1 in pro-NHEJ activity has been greatly expanded upon through a number of recent reports showing that it localizes the Shieldin protein complex, directly inhibiting exo-nucleolytic activities and long-range resection (19-25) as well as the fill-in polymerase-α that counteracts resection (22). Consistent with our data demonstrating that 53BP1 localization at DSBs in euchromatic regions requires ZEB1, reduction or loss of ZEB1 results in an increase in HR-promoting resection (Figure 3C).

Recombination between coding gene segments in V[D]J distal end-joining requires DNA-PK, Artemis and the cNHEJ repair machinery (58). As localization to ZEB1 at a LASER-induced lesion is DNA-PK dependent, our results demonstrating that ZEB1 KD significantly reduces V[D]J-mediated fluorescence output supports its role in this DNA-PK-dependent repair pathway. This result is further consistent with the strong T-cell phenotype observed in a ZEB1 loss-of-function mouse model (59). These animals have an extreme hypocellularity of the thymus with a 99% reduction in intra-thymic c-kit^+^ T-precursor cells. All thymic T-cell developmental stages– double-negative (DN), double-positive (DP) and single-positive (SP) cells–are reduced in number, but compared to wild-type controls, the ratio of the KO DN population to the other stages is higher. This has been ascribed to an up-regulation of the normally ZEB1-repressed integrin VLA-4 in pre-T cells, which inhibits their ability to migrate from the bone marrow to populate the thymus (59, 67). V[D]J recombination is initiated in the DN population, and cells having undergone productive (positively-selected) rearrangements are stimulated to proliferate, differentiate and progress into the DP stage. DN cells which fail in this regard–via recombination defects or other-wise–are blocked in their proliferative and differentiative capacity and eliminated. The phenotype of this ZEB1 loss-of-function mouse is consistent, therefore, with a defect in V[D]J recombination of the T cell receptor (either in combination with or distinct from the up-regulation of VLA-4).

In summary, we have demonstrated that ZEB1 is required for localizing 53BP1, promoting NHEJ, and inhibiting HR at DSBs generated in transcriptionally active, euchromatic regions of the genome, and that these activities are DNA-PK-dependent and ATM-independent. These findings, together with our observations on the effects of ZEB1 depletion on experimental analogues of physiological DSB repair, make a strong case for a role for ZEB1 as a required component of cNHEJ, a repair pathway which is utilized in both V[D]J and CSR (53, 54, 56, 58, 68), as well as in the majority of “clean” enzymatic DSBs (36).

## Materials and Methods

### Cell Culture

HEK293T, U2OS, M059K and M059J cell lines were obtained from ATCC. The U2OS-based DIvA cell line was a generous gift from G. Lagube (CNRS, Toulouse). The U2OS-based EJ-DR cell line was a generous gift from S. Powell and R. Bindra (Sloan Kettering). HEK 293T, I-SceI Knock-In 293T, 293T ZEB1 KO, U2OS and U2OS ZEB1 KO cells were cultured in DMEM (+ 4.5g/L glucose)/10% FCS/1X Penn-Strep. DiVA and ZEB1 KO DIvA cells were cultured in DMEM (+ 4.5g/L glucose)/10% FCS/1mM NaPyruvate/1X Penn-Strep. EJ-DR cells were cultured in DMEM (+ 4.5g/L glucose)/10% Tet-minus FCS/1X Penn-Strep. The CH12.F3 mouse B lymphocyte cell line (a generous gift from T. Honjo (Kyoto Univ.)) was cultured in RPMI/10% FCS/5% NCTC-109 (Invitrogen)/1X Penn-Strep/50μM αMeOH. MOJ/MOK human GBM cells were cultured in DMEM:F12/10% FCS/1X Penn-Strep. All cells were cultured in a humidified atmosphere of 5% CO_2_ at 37°C, and were verified mycoplasma negative.

### Plasmid Constructs

Full-length (fl) ZEB1 cDNA, (variant 2, NCBI Ref. Seq.: NM_030751.5) was amplified (see table for primers), with Bam HI (5’) and Not I (3’) overhangs and cloned via Gibson Assembly (GA) into the CMV-promoter driven expression vector pcDNA3.1, yielding pcDNA3.1flZEB1. A 183bp region encompassing the homeodomain was deleted using Q5 Polymerase mutagenesis (NEB Q5 Site-Directed Mutagenesis Kit; see table for primer sequences), yielding pcDNA3.1ΔHD. flZEB1 (var. 2) was amplified with Hind III (5’) and Bam HI (3’) overhangs (see table for primer sequences) and fused in frame at its carboxyl end via GA to eGFP in the vector eGFP-N3 (CLONTECH), to generate flZEB1-GFP. 53BP1-RFP was gift from S. Jackson, (Cambridge).

### CRISPR KO of ZEB1

293T, U2OS, and DIvA-AID cells were each transiently co-transfected (3μl linear PEI to 1μg total DNA) with the Cas9 expression vector Sp5 (Sant Cruz Biotech.) and an sgRNA-expression vector harboring a ZEB1-targeting sequence (see table). After 5h media was replaced and 48h later cells were subjected to Puromycin selection (1μg/ml for 293T, 1.5 μg/ml for U2OS and DIvA cells) for an additional 48h. Cells were then detached and serially-diluted in regular media (w/o puromycin) onto 15cm T.C. dishes to achieve well-separated single-cell-derived colonies. Colonies were picked, expanded and screened for disruption of the ZEB1 gene via PCR (see table) as well as for protein levels via WB.

### Endogenous Co-IP

U2OS cells subjected to 8Gy or no IR control were allowed to recover for 30’ at 37°, placed on ice and washed 1X with ice-cold PBS. Cells were scraped, pelleted at 4°, transferred to microfuge tubes and lysed in Buffer A 20mM Hepes (7.4), 150mM NaCl_2_, 50μg/ml EtBr, 0.2% NP-40, 0.2mM EDTA, 5mM MgCl_2_, 1mM DTT, 50U Benzonase/ml, 10% glycerol, protease and phosphatase inhibitors (Roche), 1mM PMSF, for 30 minutes at 4°. After sonication (30”on/30” off, 10’ total time) in a Bioruptor (Diagenode) to disrupt aggregates, lysate was spun at 20,000Xg @4°, 15’, and protein concentration was measured in the saved supernatant (BCA assay, Pierce). After pre-clearing the lysate with protein G sepharose for 2hr at 4°, 500 μg of protein was subjected to IP o/n at 4° with rotation using 5 μg of specified antibody, followed by protein G sepharose for an additional 2hr. Beads were washed 3X with 1ml Buffer A (no EtBr), then 2X in Buffer A with 300mM NaCl. Protein complexes were eluted in 2X SDS sample buffer, and subjected to standard WB on nitrocellulose membranes using the indicated antibodies (listed in Table).

### Survival Assays

Clonogenic survival assays were carried out as described (35).

### Chromosomal aberrations

Chromosomal aberrations at metaphase were analyzed as previously described (34, 69)

### EJ-DR-based DSB Repair Pathway Choice Assay

I-SceI-mediated DSBs were induced in the EJ-DR cell line as described (40,41). Briefly, at the appropriate time after administration of RNAi and rescue constructs, cells were rinsed 1X in 37° PBS, and replenished with complete media (with 10% Tet-minus FCS) containing 1μM Sheild1 (Aobious) and 0.1 μM triamcinolone acteonide (Cayman). After 24h, cells were again rinsed with PBS, replenished with complete media and incubated at 37° for an additional 96 hrs to allow expression and maturation of fluorescence proteins. Cells were dissociated from the culture dish using Accutase (Innovative Cell Technologies), diluted directly in PBS/1% FCS and analyzed by FLOW.

### Analysis of CSR

Wild-type or ZEB KD CH12.F3 B lymphoma cells induced to undergo CSR from IgM to IgA as described (60). Cells were split at a density of 5 (10)^5^/ml and the next day, stimulated to undergo CSR as described, by addition of 1μg/ml anti-murine CD40 Ab; BD Biosciences), 5 ng/ml recombinant murine IL-4 (R&D Systems), and 5 ng/ml recombinant human TGF-β (R&D Systems). After 72 h, cells were collected and stained on ice with Alexa-fluor 488-coupled anti-IgA Ab (Southern Biotech) and 7AAD (Roche) then analyzed by FLOW.

### Analysis of V[D]J recombination

The GFPi reporter and CMV-RAG1, CMV-RAG2 expression constructs were a generous gift of J. Demengeot (Inst. G. de Ciencia, Portugal). Recombination Activation Gene (RAG)-mediated induction of DSBs, re-orientation and subsequent expression of GFP (GFPi) cDNA was carried out as described (61). 70% confluent wt or ZEB1 KO 293T cells were co-transfected (linear PEI) with the GFPi reporter alone (background) or in combination with CMV-driven RAG1 and RAG2 expression constructs, at a ratio of 1μgGFPi : 0.32 μgRAG1 : 0.28 μgRAG2. The following day, media was changed and after an additional 48h, cells were harvested and analyzed by FLOW.

### RNAi

Knock-down of the factors in Figs 1C,D, 2C and 3B were carried out using 100 pmol Smart-pool SiRNAs (ZEB1, MRE11, Dharmacon). Knock-down of factors in Fig 3A was carried out using 100 pmol Smart-pool SiRNAs (53BP1, BRCA1, NT control, Dharmacon); or, in the case of ZEB1, 25 pmol siRNA targeting the ZEB1 3’ UTR (IDT; list of sequences below). One million cells were electroporated (Nucleofector, Lonza), and after 48h, ZEB1 KD EJ-DR cells were rescued via transfection (3:1 ratio of PEI to total DNA) with either empty vector or indicated expression constructs in Fig 3A, and cultured for an additional 48h before induction of DSBs. KD of ZEB1 in the mouse B lymphoma cell line CH12.F3 was carried out using 25 pmol siRNA targeting the ZEB1 3’ UTR, as above.

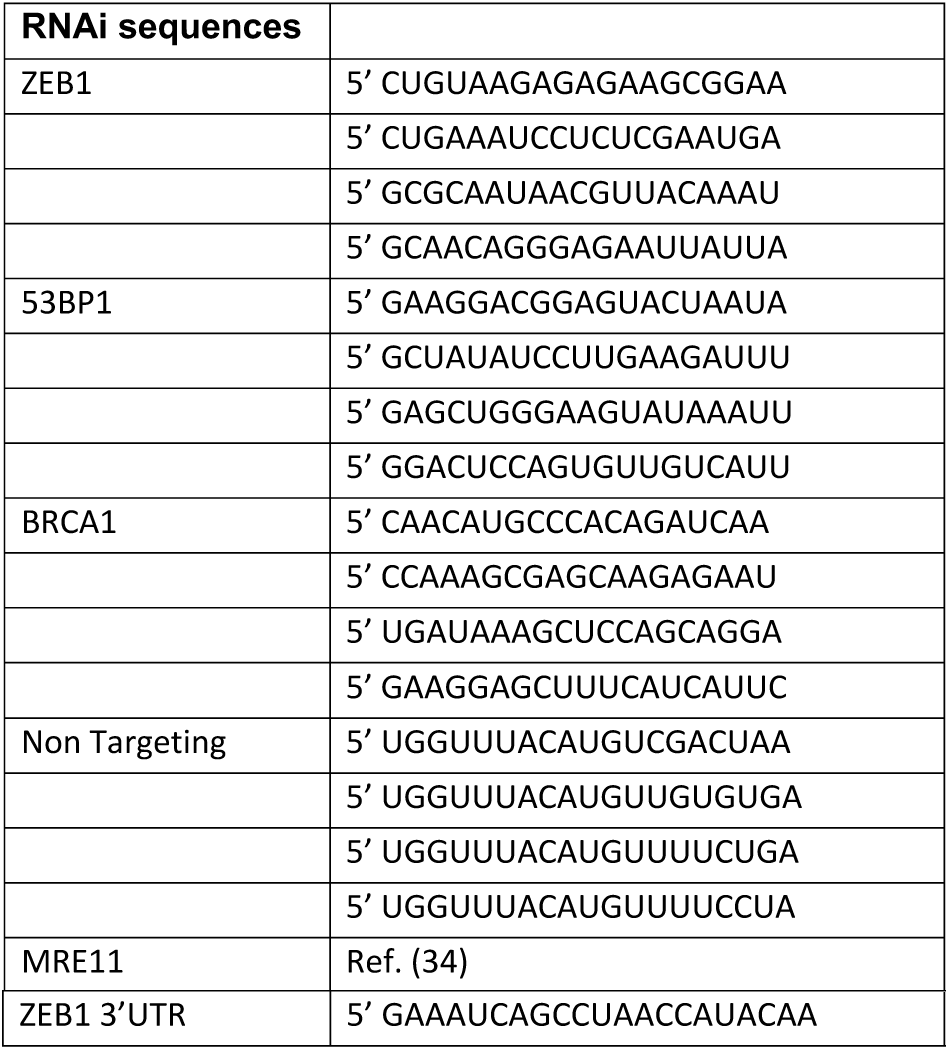

### ChIP

In the case of 293T cells (I-SceI-induced DSBs), ChIP was carried out as described (SMARCAD1). For the U2OS-based DIvA system, the protocol of Nelson, et.al. (Nat Protoc. V1 2006, p79) was followed with minor variations. After fixing in 1% formaldehyde for 10’ and Glycine quenching, cells were rinsed with ice-cold PBS, scraped, pelleted, and resuspended at 50μl/10^6^ cells in Lysis buffer (50mM Tris pH8.0,150mM NaCl, 5mM EDTA, 1%Triton X-100, 0.1% Na Deoxycholate, 0.5% SDS, protease inhibitors and phosphatase inhibitors (Roche), 1mM PMSF). Chromatin was sonicated to an avg size of 250-750bp and 5% percent of the cleared lysate was removed for input. The remainder was diluted 5-fold in Chip dilution Buffer (50mM Tris-Cl pH8.0,150mM NaCl, 2mM EDTA, 1.2%Triton X-100, protease inhibitors and phosphatase inhibitors, 1mM PMSF). After addition of Abs (or isotype-specific IgG; see table), samples were rotated at 4° o/n, followed by addition of biotinylated secondary Ab, rotation for 1h, 4°, and then with pre-blocked streptavidin-coupled magnetic beads, and further rotation for 2h, 4°. Beads were washed 3X with ice-cold (20mM Tris-Cl pH8.0,150mM NaCl, 2mM EDTA, 0.1% SDS, 1% Triton X-100) and then once with ice-cold (20mM Tris-Cl pH8.0, 500mM NaCl, 2mM EDTA, 0.1% SDS, 1% Triton X-100). Following the Chelex-100 isolation step, DNA was purified using a standard PCR-clean-up column (QiAGEN). 2 μl of purified ChIPed DNA was used in the subsequent qPCR reaction.

### qPCR

Real-time qPCR analysis was carried out using iTaq Universal SYBR Green Supermix (Bio-Rad) for 39 cycles on a CFX96RealTime PCR machine (Bio-Rad). Data were calculated using the ΔCt method and presented as percent of input. All data shown are the results of at least three independent ChIP experiments with qPCR performed in triplicate and results averaged with SEM.

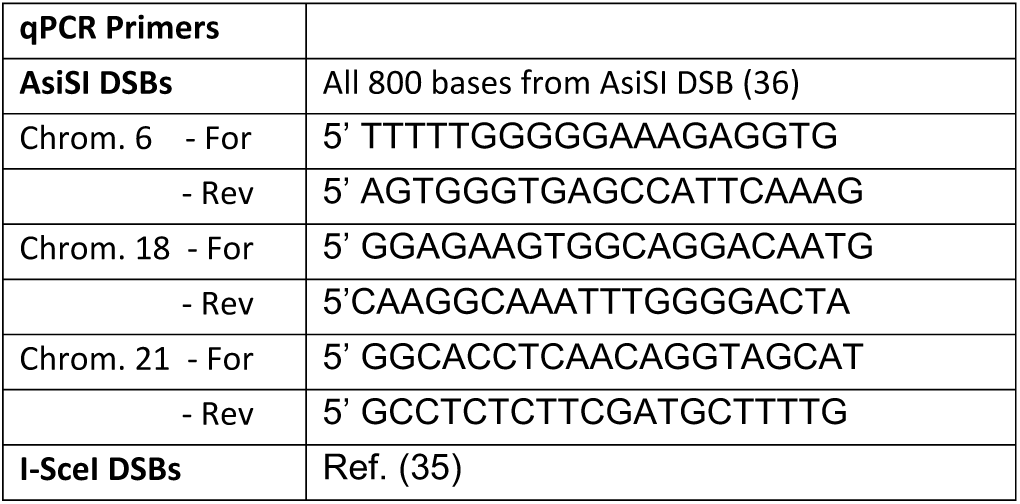

### DNA End-Resection Assays

Isolation of genomic DNA, o/n digestion with indicated restriction enzymes, and subsequent qPCR analysis was performed as described (X). The extent of resection at an AsiSI-induced DSB was quantitated in asynchronous or synchronized DIvA cells as previously described (34). Synchronization was achieved using double thymidine block (Cohen ref), with G1 and G2 cells collected 15hr and 7hr post-release from the second block, respectively. After o/n digestion with Bsr GI, genomic DNA was purified and ssDNA near the AsiSI-cut site on Chromosome 1 measured by qPCR, using the Taq-Man assay (see ref. 34 (SMARCAD1) for primers).

### Immunofluorescence Staining

Co-immunofluorescence staining of 53BP1 and ZEB1 (Fig 1B) was carried out as described (34). Immuno-staining for γH2AX was carried out as follows. Wild-type or ZEB1 KO 293T cells were grown on poly-lysine-coated coverslips, and, at given time points, IR-treated (or no control) cells were gently rinsed in 1X PBS, fixed in freshly-made 4% paraformaldehyde for 15’, permeabilized in 0.1% Triton X-100/PBS, 5’, and blocked for 1h in 1%BSA/0.05%TritonX-100/PBS (all at RT), then stained o/n at 4° with an anti-γH2A.X-Alexafluor488 Ab (see Table) diluted 1:2500 in blocking buffer. After washing 3X, 5’ in 0.05%Triton-X100/PBS at RT, with gentle rocking, coverslips were mounted and imaged on a Zeiss LSM-700 inverted microscope using a 20X objective.

### Live Cell Imaging

U2OS or M059J/K cells were plated onto 35mm glass bottomed dishes (Ibidi) and 24h later transfected with the indicated fluorescence reporter constructs (ratio of 3:1 linear PEI to DNA), and cells were imaged 48h later. Initial plating numbers were calculated so that cells were growing exponentially on day of imaging. One hour before imaging, media was exchanged for its identical phenol red-minus counterpart (DMEM for U2OS; DMEM:F12 for M059J/K GBM cells). To inhibit DNA-PK activity in U2OS cells, NU7441 (20nM final concentration) or DMSO vehicle was added at this time. Laser ablation and live cell imaging were carried out on a Zeiss LSM 780 laser scanning confocal microscope with an integrated IR Chameleon Vision-II, ultrafast Ti:Sapphire laser. Dishes were imaged on a heated stage under sparged, 5% CO_2_. To minimize confounding cellular damage, the focal plane was positioned at the center of each nucleus and a thin rectangular region of interest was drawn that traversed the nucleus while avoiding the nuclear membrane. Images were recorded through a 40X oil immersion objective using continuous capture as follows. Two images (1.94” total time per image to capture both color channels) were recorded and the laser (tuned to 780nm, 10% power, for 3 iterations that combined totaled 12μsec.) was triggered at the beginning of the third image capture.

### Statistics

Bar graphs represent the mean ± SEM of at least three independent experiments. The unpaired two-tailed Student’s t test was used to calculate statistical significance, and a P < 0.05 value was considered significant.

## List of Primers

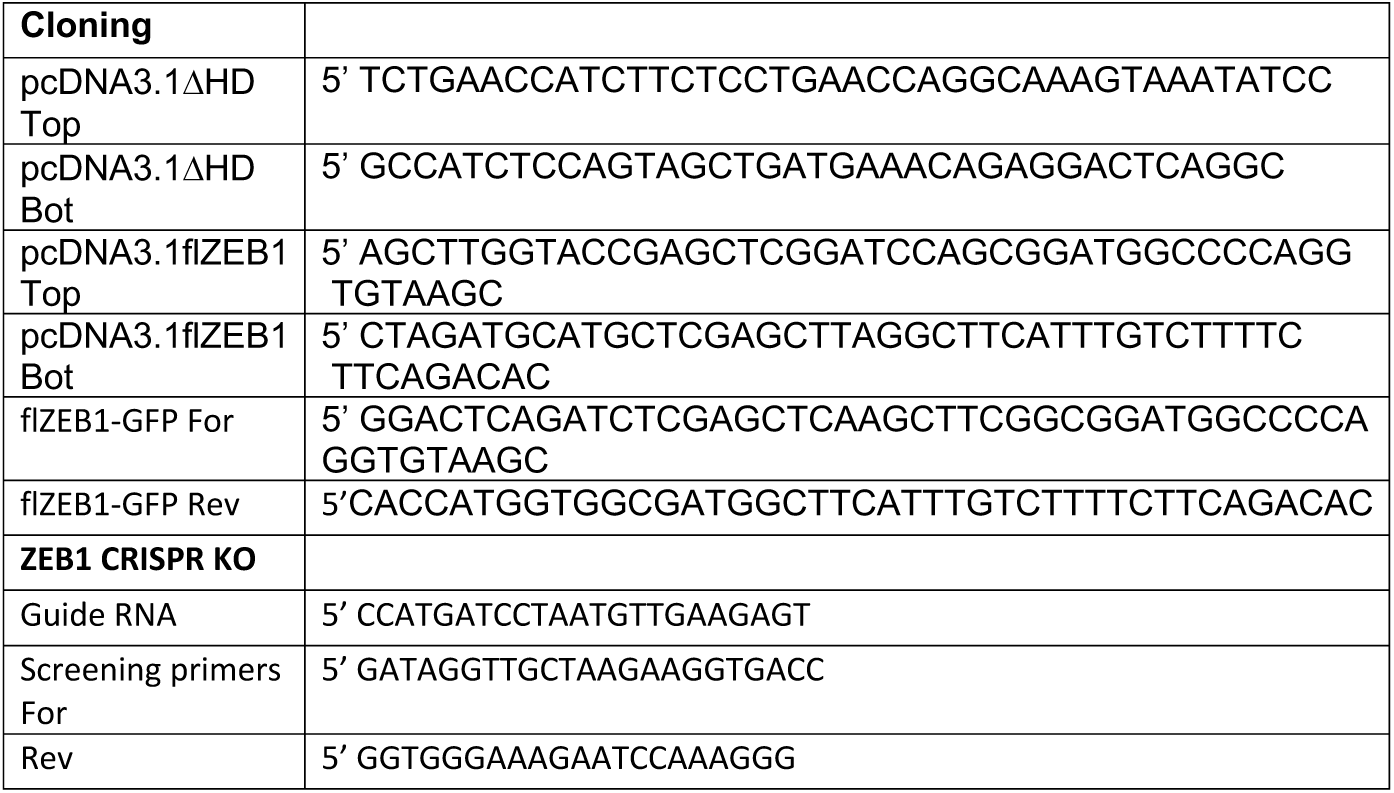

## Antibodies Used in This Study (all human)

|  |  |  |
| --- | --- | --- |
| ZEB1 | Bethyl | A301-921A (IF, WB) |
|  | Invitrogen | 21544-1-AP (ChIP) |
| 53BP1 | Millipore | MAB3802 (ChIP, IF, WB) |
| α-Tubulin | SIGMA | T9026 (WB) |
| γH2A.X Alexa Fluor-488 | ThermoFisher | 53-9865-82 (IF) |
| HDAC1 | Cell Signaling | 2062 (WB) |
| HDAC2 | Cell Signaling | 2545 (WB) |
| H3K9Me3 | abcam | ab8898 (ChIP) |
| PRDM2 | ThermoFisher | MA5-11144 (ChIP) |
| MacroH2A1.1 | abcam | Ab37264 (ChIP) |

## Key Reagents/Supplies

| Reagent | Source |
| --- | --- |
| 4-OHT | SIGMA |
| Sheild1 | Aobious |
| Triamcinolone acteonide | Cayman |
| Accutase | Innovative Cell Technologies |
| CD40 Ab | BD Biosciences |
| Recombinant IL-4 | R&D Systems |
| Recombinant human TGF-b | R&D Systems |
| Alexa-fluor 488-coupled anti-IgA Ab | Southern Biotech |
| 7AAD | Roche |
| Linear PEI 25K | Polysciences |
| NU7441 | Tocris |
| KU5993 | Tocris |
| Benzonase | SIGMA |
| Chelex-100 | Bio-Rad |
| 35mm Glass-bottomed dishes | Ibidi |

## Acknowledgments

The authors thank R. Bindra, S. Powell, G. Lagube, T. Honjo, S. Jackson, G.M. Anderson, and J. Demengeot for cell lines and plasmid constructs, and M. Huang and L. Lum for help with FLOW. We thank R. Cao and A. Periasamy for help with live cell image capture, which was acquired using the UVA Keck Center Zeiss 780 Confocal microscopy system (NIH OD016446). This work was supported by The Charles Burnett III Fund.

